# Household socio-economic status and likelihood of HIV infection among under five-year children in Muheza district, north-eastern Tanzania

**DOI:** 10.1101/661801

**Authors:** Veneranda M. Bwana, Edgar Simulundu, Leonard E.G. Mboera, Sayoki G. Mfinanga, Charles Michelo

## Abstract

**Background:** There are evidences of the association between socio-economic factors and HIV prevalence in Sub-Saharan Africa. However, there is dearth of information on such relationship in Tanzania. Here, we present data on the relationship between household’s socio-economic factors and HIV prevalence among under five-year children in Muheza district, Tanzania.

**Methods:** We conducted a facility-based study from June 2015 to June 2016 in which we enrolled under five-year children born to HIV positive mothers. Information on HIV status of the child and socio-demographic characteristic of the head of the household was collected using a structured questionnaire. Data analysis was done using STATA version 13.0.

**Results:** A total of 576 mothers/guardians were interviewed each with respective HIV exposed under five-year child. Children who belonged to a head of household with at least a high education level (AOR= 0.4, 95% CI 0.2-0.8) and living in a relatively wealthy household (AOR = 0.5, 95% CI 0.2-0.9) was associated with reduced odds of HIV infection among children. Univariate analysis revealed that the odds of HIV infection was three-fold (COR = 2.9, 95% CI 1.2-7.0) higher among children living in rural than in urban areas. The heads of household living in rural areas (AOR=0.3 95% CI 0.1-0.9) had low education level compared to those living in urban areas.

**Conclusion:** Children who belong to the head of households with high educational level, high household wealth were associated with reduced likelihood of HIV infection in Tanzania. Children living in rural areas had increased likelihood of acquiring HIV infection. These findings stress the need to focus on improving education status of the population and economically disadvantaged populations as a strategy for HIV prevention and control measures.

## Introduction

By the end of 2016, Sub-Saharan Africa had the highest burden of paediatric HIV with over 90% of the estimated 2.1 million children living with HIV worldwide [2]. Tanzania is one of the countries in Sub-Saharan Africa most devastated by this global pandemic. In 2016, estimated 1.4 million people in Tanzania were living with HIV and 18% of these infections were due to mother to child transmission (MTCT) [3]. Maternal immunological status with a Cluster of Differentiation 4 (CD4) cell count less than 200 cell per mm^3^ near delivery, high maternal viral load, advanced stage of maternal HIV infection have been observed to increase the risk of MTCT [4, 5]. These risk factors operates through socio-economic statuses such as occupation, marital status, education and income to bring a health outcome among children [6, 7].

Studies have indicated that higher education attainment is a proxy for higher socio-economic status which is an indication of increased wealth [8]. It has been argued that in early HIV epidemic, higher education achievement was associated with higher wealth, and increased mobility that potentially increases exposure to HIV infection [9]. However, as the epidemic matures the pattern shifts towards reduced risk among the wealthy and higher educated individuals [10-12]. Whilst HIV has been classified as the disease of the poor, the causality of which one cause the other between poverty and HIV is complex and challenging [13, 14]. Over the recent years, the burden of HIV has been reported to be greater among the poorer and least educated populations [11, 15, 16]. This is likely to be due to their engagement in risky sexual behaviours and transactional sexual relationships and limited access to health facilities [17].

According to the World Bank report, it has been observed that improved education leads to poverty reduction, boosts economic growth and increases income [18]. Countries with high HIV/AIDS burden are continuously stressing the need to put the goal for universal education for all as an important Sustainable Development Goal as well as a strategy for combating the HIV/AIDS epidemic [19-22]. In countries with mature HIV epidemics, whereby HIV/AIDS education programmes have been introduced, high education attainment appears to become protective against HIV [11, 16]. Higher education levels attainment has been associated with decreased risk of HIV infection especially among younger people in Ethiopia, Zambia, Uganda and Tanzania [10, 23-25]. This association has been attributed, in part to increased awareness, adoption of safe behavioral practices and prompt access to preventive and curative health services by the more educated individuals [13]. Several existing evidence in support of the positive impact of education on better health outcomes are available [11, 14, 16, 26, 27], most of them suggest that education attainment imparts similar influences on HIV burden, reduced child mortality and increased survival.

Little is known about the effect of head of households’ socio-economic (SES) on HIV prevalence of their children in Tanzania. It is important that this relationship is investigated in order to understand factors influencing paediatric HIV. This study was therefore carried out to determine the relationship between household SES and HIV prevalence among under five-year children in Muheza district, Tanzania.

## Methods

### Study area and population

This facility-based study was carried out from June 2015 to June 2016 in Muheza district, in north-eastern Tanzania (4°, 45’S; 39°00’E). The study population involved mothers/guardians with their respective HIV exposed under five-year children. Using a multistage sampling approach, we enrolled a random sample of mother/guardian child pairs. More details of this study have been described elsewhere [28]. More than 79% of households in the district are involved in agriculture as the main source of livelihood and income. The remaining proportion include petty traders, fisherman and livestock keepers. The district has 116 primary schools and 31 secondary schools [29]. The district has a total of 46 health facilities which include one hospital, four health centers and 41 dispensaries.

### Data collection

Socio-demographic factors of the mother/guardian-child pairs (education, age, sex, marital status, distance from a health facility, occupation, residence; head of household (age, sex, education, size of household, proxy variables for household wealth) were collected using a structured questionnaire. The household’s wealth was also assessed based on four key questions; namely primary house construction materials, main source of fuel used for cooking, ownership of land and presence/absence of electricity.

Information on the HIV status of the children was extracted from data-base available at the district hospital. Data on education level was based on the Tanzanian system whereby, individuals attend 7 years of primary education, 4 years of secondary education (ordinary level), and 2 years of advanced secondary education. After ordinary secondary education or advanced secondary education, individuals meet criteria for college education or qualify for University education after advanced level of secondary education.

### Data management and analysis

Data analysis was performed using Stata software version 13. For the child HIV status was categorized into a binary outcome variable as “HIV positive” and “HIV negative”. Education level attainment was categorized into a binary outcome (‘low education’ or ‘high education’), measured as the number of formal school years attended by the respondent. Low education level was recorded as 0-6 years schooling and high education as 7+ years schooling. The household wealth was categorized into a binary variable ‘high’ or ‘low’. Size of the household was categorized into two: less or equal to seven people and more than seven people living in the same household. Multiple logistic regression was done by including variables from univariate analysis with P-value of ≤0.2. Multiple logistic regression analyses were used to examine the associations between various socio-economic factors of the head of household and the child’s HIV infection status. Backward Logistic Regression method was employed by removing variables with highest P-value until the remaining variables in the final model had a P-values ≤ 0.05 and adjusted odd ratios (AOR) at 95% confidence interval. Crude odd ratios (COR) with corresponding P-values were also presented.

### Ethical considerations

Ethical approval was obtained from Medical Research Coordinating Committee of the National Institute for Medical Research in Tanzania with reference number NIMR/HQ/R.8a/Vol. IX/1978. Permission to conduct this study was granted by Muheza District Council Authority. Written informed consent was obtained from each mother/guardian before recruitment.

## Results

### Socio-demographic characteristics of the respondents

A total of 576 mothers/guardians each with HIV exposed under five-year child were interviewed. Out of 576 under five-year children, 281 (48.8%) were males and 295 (51.2%) were females. A total of 379 (65.8%) under five-year children were aged ≤ 24 months and 197 (34.2%) aged 25-59 months. More than half (n=309) of under five-year children were livingfar (more than 30 minutes walk by foot) from a health facility. A total of 61 under five-year children were confirmed to be HIV positive whereby 75.4% (46) of them, belonged to the head of households who had completed primary (7 years) level of education (Figure 1). The education level of the heads of household were: 58 (10.1%) had less than 7 years of education, 518 (89.9%) had 7+ years of education. Majority (82.1%, n=473) of the heads of household had completed primary (7 years) level education. Forty-two (7.3%), 3 (0.5%), had 8-11 years and 12+ years of schooling respectively. A total of 445 (73%) household heads were living in rural areas. Households with low wealth were 441 (77%) and those with high wealth were 135 (23%).

**Figure 1:**
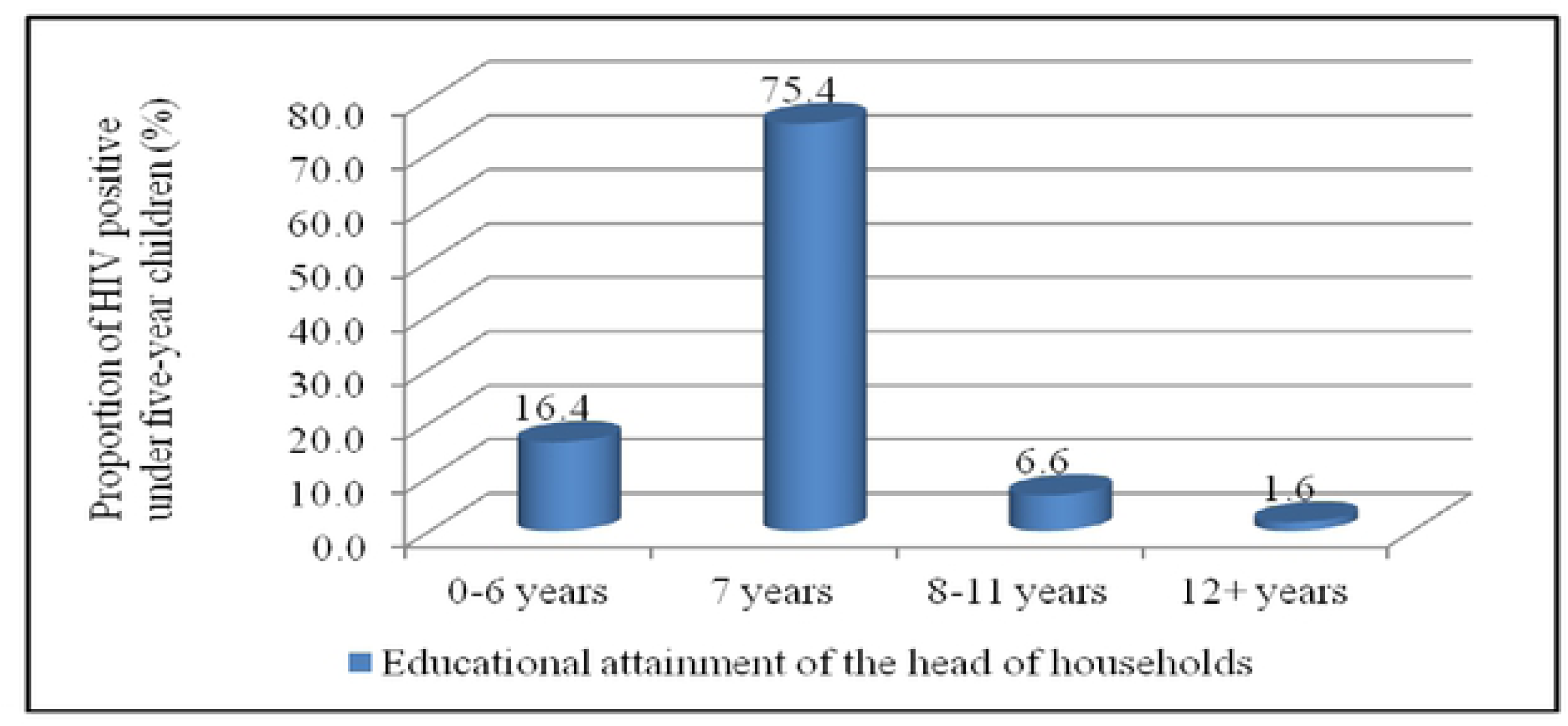
Proportion of HIV positive under five-year children by educational attainment of their head of households.

### Socio-economic factors associated with HIV infection among under-five year children

In multiple logistic regression, higher odds of HIV infection was observed among children aged more than two years (AOR=3.0, 95% CI 1.5-5.9) than in younger children. Likewise, higher likelihood of HIV infection was 2.4 (AOR= 2.4 (1.2-5.0) times higher among those who lived far from a health facility as compared to their counterparts. The odds of HIV infection in children was lower (AOR= 0.4, 95% CI 0.2-0.8) among those with a head of household who had attained high education level and in those living in households with high wealth (AOR = 0.5, 95% CI 0.2-0.9) (Table 1). At univariate analysis, the odds of HIV infection was 3 times higher among children living in rural areas than urban areas. Guardian’s marital status, occupation and education did not show any association with HIV infection among under five-year children.

**Table 1:** Household’s socio-economic factors associated with HIV infection among under five-year children.

| **Variables** | **Proportion (%)** | **COR**<br>**(95% CI)** | **\$P value** | **AOR**<br>**(95%CI)** | **\*P value** |
| --- | --- | --- | --- | --- | --- |
| Child age (months) |  |  |  |  |  |
| ≤24 | 65.8 | 1 |  | 1 |  |
| 25-59 | 34.2 | 5.1 (2.9-9.1) | < 0.001 | 3.0 (1.5-5.9) | 0.002 |
| Residence |  |  |  |  |  |
| Urban | 22.7 | 1 |  | 1 |  |
| Rural | 77.3 | 2.9 (1.2-7.0) | 0.02 | 1.5 (0.6-4.0) | 0.4 |
| Distance to health facility |  |  |  |  |  |
| Near ($\leq 30$ minutes) | 46.4 | 1 | | 1 | |
| Far ($> 30$ minutes) | 53.6 | 3.2 (1.7-6.0) | $< 0.001$ | 2.4 (1.2-5.0) | 0.02 |
| Head of household education |  |  |  |  |  |
| Low (0-6 years schooling) | 10.1 | 1 |  | 1 |  |
| High (7+ years schooling) | 89.9 | 0.5 (0.3-1.1) | 0.08 | 0.4 (0.2-0.8) | 0.02 |
| Household wealth |  |  |  |  |  |
| Low | 77 | 1 |  | 1 |  |
| High | 23 | 0.8 (0.4-1.6) | 0.2 | 0.5 (0.2-0.9) | 0.03 |

**Notes:** (a) Total sample size, N = 576 (b) \*P values of AOR; §P values of COR (c) COR = Crude odd ratio; AOR = Adjusted odd ratio; CI = Confidence interval

### Head of household’s education

Results of the multiple logistic regression model showed significant association between level of education, and gender, size of the household, wealth. Males who were heads of household (AOR=2.0, 95% CI 1.1-3.5) had high education level attainment compared to their counterpart females. The households with high wealth were significantly associated with high education level of the heads of household (AOR =1.5, 95% CI 1.3-2.0). Heads of household living in rural areas (AOR=0.3 95% CI 0.1-0.9) had low education level compared to those living in urban areas. In the household with more than seven people living in the same house (AOR=0.5, 95% CI 0.2-0.9), was significantly associated with low education level of the head of household (Table 2).

**Table 2:** Factors associated with high education level attainment of the heads of household.

| Variables | Proportion (%) | COR<br>(95%CI) | \$P value | AOR<br>(95%CI) | *P value |
| --- | --- | --- | --- | --- | --- |
| Head of household sex |  |  |  |  |  |
| Female | 29.9 | 1 |  | 1 |  |
| Male | 70.1 | 2.2 (1.3-3.9) | 0.004 | 2.0 (1.1-3.5) | 0.02 |
| Residence |  |  |  |  |  |
| Urban | 22.7 | 1 |  | 1 |  |
| Rural | 77.3 | 0.2 (0.1-0.6) | 0.01 | 0.3 (0.1-0.9) | 0.03 |
| Size of household |  |  |  |  |  |
| ≤ 7 people | 90.1 | 1 |  | 1 |  |
| > 7 people | 9.9 | 0.4 (0.2-0.7) | 0.01 | 0.5 (0.2-0.9) | 0.04 |
| Household wealth |  |  |  |  |  |
| Low | 77 | 1 |  | 1 |  |
| High | 23 | 1.4 (1.3-1.8) | 0.01 | 1.5 (1.3-2.0) | 0.02 |

**Notes:** (a) Total sample size, N = 576 (b) \*P values of AOR; \$P values of COR (c) COR = Crude odd ratio; AOR = Adjusted odd ratio; CI = Confidence interval

## Discussion

Our findings indicate that high education level of the heads of household and high household wealth have higher probability of reduced odds of HIV infection among under five-year children. The likelihood of acquiring HIV infection is higher among under five-year children located in rural than in urban areas. Households with higher wealth were significantly associated with high education attainment of the heads of household. Moreover, the heads of household located in rural areas had low education level attainment compared to those in urban areas.

Education is one of the most important component of social and economic development [30]. Education has positive spill-over effects whereby all people in the society will benefit even if only few of them have received education [13]. The observed reduction on the likelihood of HIV infection among under five-year children in the households with heads with higher education level in this study could be due to adoption of preventive strategies on HIV/AIDS among higher educated individuals [13]. A study in Zambia has reported that the number of years one spent in school have an impact on the knowledge and ability to follow prevention strategies against HIV/AIDS transmission, that influence better health outcomes and child survival in the community [11, 16, 31]. The assumption that effects of educational attainment of the head of household on the likelihood of acquiring of HIV infection in children is likely to be exerted through mediator factors such as the power of decision to send the child to clinic to access interventions [32, 33]. It has been argued that, a set of proximate determinants can be influenced by changes in socio-economic determinants or by interventions which have a direct effect on biological mechanisms to influence the risk of morbidity and mortality among children (34). This implies that educational attainment in itself cannot, however, be isolated from other socio-economic factors as the grounds of HIV risk reduction.

Moreover, levels of knowledge, literacy and education tend to be lower among the poor and individuals living in rural areas which greatly influence household decisions with regard to the proximate determinants of health [18, 35]. However, educational opportunities might be quite limited in rural areas with poor road infrastructures that make the formal school system difficult to enter. A previous study in Tanzania, demonstrated increased HIV prevalence in rural population with less educated individuals at an increased risks [10]. Studies conducted in Sub Saharan Africa have reported that children living in rural areas are more prone to home deliveries that predispose them to HIV infection [36-38]. This could be attributed to inaccessibility to health promotion messages regarding HIV/AIDS prevention in rural compared to urban settings or women’s participation in household decision making to access health related information and services in those male-headed household.

Higher educational achievement associated with higher household wealth observed in this study has also been reported elsewhere [14, 35, 39]. More-educated individuals are most likely to have more income and thus more control over their living [9]. They are likely to place higher value on future endeavors and thus be more motivated to adopt preventative measures particularly against infectious disease risks [40]. In addition, the interaction effects between education and wealth could be very positive; when individuals have resources and the ability to use those resources, they can act on safeguarding their health seeking behavioral patterns [41]. This suggest that a convincing interaction do exist between wealth and education, as well as between education and the risk of acquiring diseases among family members living in the same household. Moreover, higher income at household level is interrelated with higher educational attainment and better health outcomes which is attributed, in part to more frequent and more intensive use of health services at both private and public health sectors [42]. A recent study in South Africa reported decreased risk of HIV infection among individuals living in less poor household and having tertiary education [43]. This indicates that strategies that enhance economic empowerment and provision of quality education to every individuals is highly recommended in order to bring positive health outcomes in the society.

The study has limitations including the possibility that our results could have been affected by unmeasured confounders, which cannot be eliminated. Since some variables may not fully be adjusted during analysis, especially when dealing with proxy variables for household wealth which might be subjected to misclassification. Selection bias, as only those who have attended at the health facility got higher chance to be selected. These individuals may differ from those who were left to be unidentified in the community.

## Conclusion

High educational level attainment of the heads of household and high household wealth was associated with reduced likelihood of acquiring HIV infection among under five-year children living in the same household. Under five-year children located in rural areas had increased likelihood of acquiring HIV infection than their counterparts in urban areas. Thus, there is a need to focus on improving education and general knowledge to the whole population and economically disadvantaged populations as complementary strategies in HIV/AIDS control and prevention programmes.

## Authors’ contributions

VMB conceived the study. VMB, CCM, LEGM, SGM, ES contributed to the design of the approach. VMB analysed and interpreted the data. VMB wrote the manuscript and all authors read, contributed to, and approved the final version of the manuscript.

## Acknowledgments

We would like to acknowledge the entire administration of Muheza District Council for granting permission to to conduct this study. Thanks to Juma Mfanga, Dorothy Lema, Angelina Sengwaji, Paulo Muhusa, Herieth Nyangasa, John Kwingwa, Baltazar Nyombi, and staff of the Muheza Designated District Hospital, health centres and dispensaries for their extended cooperation during data collection. We would like to acknowledge the excellent technical and field assistance of Bernadina Ruta, Stella Kolugendo, Josephine Mhina, Irene Lazaro, Grades Stanley, Susan F. Rumisha, Fagason Mduma, Veronica Msingwa and Diana Kwetukia. Many thanks to all children and their families who agreed to participate in this study.

